# CURT1A and CURT1C mediate distinct stages of plastid conversion in *Arabidopsis*

**DOI:** 10.1101/2021.12.01.470752

**Authors:** Zizhen Liang, Wai Tsun Yeung, Keith Ka Ki Mai, Juncai Ma, Zhongyuan Liu, Yau-Lun Felix Chong, Byung-Ho Kang

## Abstract

The crystalline structure of prolamellar bodies (PLBs) and light-induced etioplasts-to-chloroplasts transformation have been investigated with electron microscopy methods. However, these studies suffer from chemical fixation artifacts and limited volumes of tomographic reconstruction. We have examined *Arabidopsis thaliana* cotyledon samples preserved by high-pressure freezing with scanning transmission electron tomography to visualize larger volumes in etioplasts and their conversion into chloroplasts. PLB tubules were arranged in a zinc blende-type lattice like carbon atoms in diamonds. Within 2 hours after illumination, the lattice collapsed from the PLB exterior and the disorganized tubules merged to form fenestrated sheets that eventually matured into lamellar thylakoids. These planar thylakoids emerging from PLBs overlapped or folded into grana stacks in PLBs’ vicinity. Since the nascent lamellae had curved membrane at their tips, we examined the localization of CURT1 proteins. *CURT1A* transcript was most abundant in de-etiolating cotyledon samples, and CURT1A concentrated at the peripheral PLB. In *curt1a* mutant etioplasts, thylakoid sheets were swollen and failed to develop stacks. In *curt1c* mutant, however, PLBs had cracks in their lattices, indicating that CURT1C contributes to cubic crystal growth under darkness. Our data provide evidence that CURT1A and CURT1C play distinct roles in the etioplast and chloroplast biogenesis.

## Introduction

Plastids exist in different forms depending on the cell type and environmental conditions (Jarvis and López-Juez, 2013; Kirchhoff, 2019). In germinating seedlings, proplastids in the cotyledon develop into chloroplasts. As chlorophyll biosynthesis is inhibited under darkness, the photosynthetic protein complexes of the thylakoid membrane are not assembled, and chloroplast biogenesis is inhibited. Instead, developmentally plastids, known as etioplasts, form (Solymosi and Schoefs, 2010). The etioplasts transform into chloroplasts once light becomes available and chlorophyll accumulates (Hernandez-Verdeja et al., 2020).

Thylakoids in etioplasts consist of semi-crystalline tubular membrane networks of prolamellar bodies (PLBs) connected by planar prothylakoids (Ryberg and Sundqvist, 1982; Rascio et al., 1984). During the light-induced etioplast-chloroplast transition, the proteins, lipids, and cofactors stored in PLBs provide building blocks for the chloroplast thylakoids (Ploscher et al., 2011; Armarego-Marriott et al., 2019; Fujii et al., 2019). The most abundant protein constituent of PLBs is light-dependent protochlorophyllide oxidoreductase (LPOR) (Blomqvist et al., 2008), which forms a helical array surrounding PLB tubules (Floris and Kuhlbrandt, 2021). LPOR is a photocatalytic enzyme that mediates the reduction of protochlorophyllide (Pchlide) into chlorophyllide to produce chlorophyll (Zhang et al., 2019). LPOR oligomerize on liposomes mimicking the PLB membrane to tubulate them *in vivo* as shown by cryo-electron microscopy (Nguyen et al., 2021). It is thought that LPOR undergoes conformational changes after the photoreduction and dissociates from the PLB membrane, resulting in the breakdown of the PLB lattice. Inactivation of *PORA*, an *Arabidopsis* gene encoding an LPOR protein, led to structural defects in PLBs and abnormal photomorphogenesis (Paddock et al., 2012).

When examined under electron microscopy, PLBs are mostly made of hexagonal lattices in which tetrahedral units repeat (Murakami et al., 1985). Small angle X-ray studies of isolated PLBs revealed that branched tubules in PLBs are packed mostly in the cubic diamond (*i*.*e*., zinc blende) symmetry (Williams et al., 1998; Selstam et al., 2007). Recent electron tomography (ET) imaging of runner bean (*Phaseolus coccineus*) indicated that the PLB lattice matched the wurtzite-type crystal symmetry (Kowalewska et. al, 2016). PLBs in which tubules deviate from the tetrahedral pattern have been reported, and the arrangement is called the “open” type (Gunning, 2001). Moreover, etiolation conditions affect the sizes and density of PLBs (Bykowski et al., 2020).

The light-triggered transformation of PLBs into grana and stroma thylakoids was first investigated with electron microscopy in the 1960s, although those early studies were based on two-dimensional electron micrographs of PLBs and thylakoids (Gunning, 1965; Henningsen and Boynton, 1974; Rascio et al., 1984; Grzyb et al., 2013). An ET analysis of the etioplast-to-chloroplast transition in runner bean cotyledons showed that PLB tubules directly coalesce into planar thylakoid elements without involvement of vesicular intermediates and that the helical arrangement of the inter-disc connections within a grana stack appears early in the granum development (Kowalewska et al., 2016). In the Kowalewska study, the etioplast volumes in the ET reconstruction were limited in the z-direction coverage, visualizing two hexagonal layers in 3D. Using serial block face-scanning electron microscopy, it was demonstrated that the conversion of PLBs into photosynthetic thylakoids in *Arabidopsis* cotyledons is within 24 hour after light illumination in concurrence with correlative proteomic and lipidomic results (Pipitone et al., 2021). The serial block face-scanning electron microscopy approach can visualize larger volumes encompassing the entire etioplasts or chloroplasts, but the resolution is poorer than ET, especially along the z-axis. In both 3D EM studies, cotyledon samples were prepared with chemical fixation, which fails to preserve intricate or short-lived structures in cells (McIntosh et al., 2005; Staehelin and Kang, 2008).

In this study, we examined etioplasts in *Arabidopsis* cotyledons grown in the dark with serial section ET. Cotyledon samples were prepared by high-pressure freezing to avoid fixation artifacts. As the stroma is heavily stained in high-pressure frozen etioplasts, we employed scanning transmission ET (STET), which enhances image contrast in tomograms from such specimens (Aoyama et al., 2008; Hohmann-Marriott et al., 2009; Murata et al., 2014; Kang, 2016). CURVATURE THYLAKOID1 (CURT1) family proteins are thylakoid membrane proteins that stabilize the sharply curved membrane at the grana margin (Armbruster et al., 2013; Pribil et al., 2014). Our time-resolved ET study of wild-type and *curt1* family mutant cotyledons indicated that grana stacks arise directly from PLBs and that CURT1A is required for the stack assembly. By contrast, CURT1C plays a role in the cubic crystal close packing for PLB biogenesis in developing etioplasts.

## Results

### Crystalline structure of the *Arabidopsis* PLBs and their light-induced degradation

To estimate the timeline of the etioplast-to-chloroplast transformation, we examined etiolated Col-0 *Arabidopsis* cotyledons at 0, 2, 4, 8, and 12 hours after light illumination (HAL). Cotyledon greening was clearly observed at 12 HAL (Fig. 1A-C), and chlorophyll autofluorescence increased during this period (Fig. S1A-E). PLBs were small spots of 1.5-2.0 µm in diameter that emitted autofluorescence in 0 HAL cotyledons (Fig. 1D). The fluorescent spots were enlarged and spread out to form ovals at 2 HAL (Fig. 1E). We monitored the degradation of PLBs with transmission electron microscopy (TEM) and STET at the five time points. Each etioplast had one or two PLBs that were linked by multiple lamellar thylakoids at 0 HAL (Fig. 1F and S1F). Lamellar thylakoids in 0 HAL etioplasts were ribosomes on their membranes, like the pre-granal thylakoids in proplastids of germinating cotyledon cells at 36 and 64 hours after imbibition reported in Liang et al. (2018).

**Figure 1.**
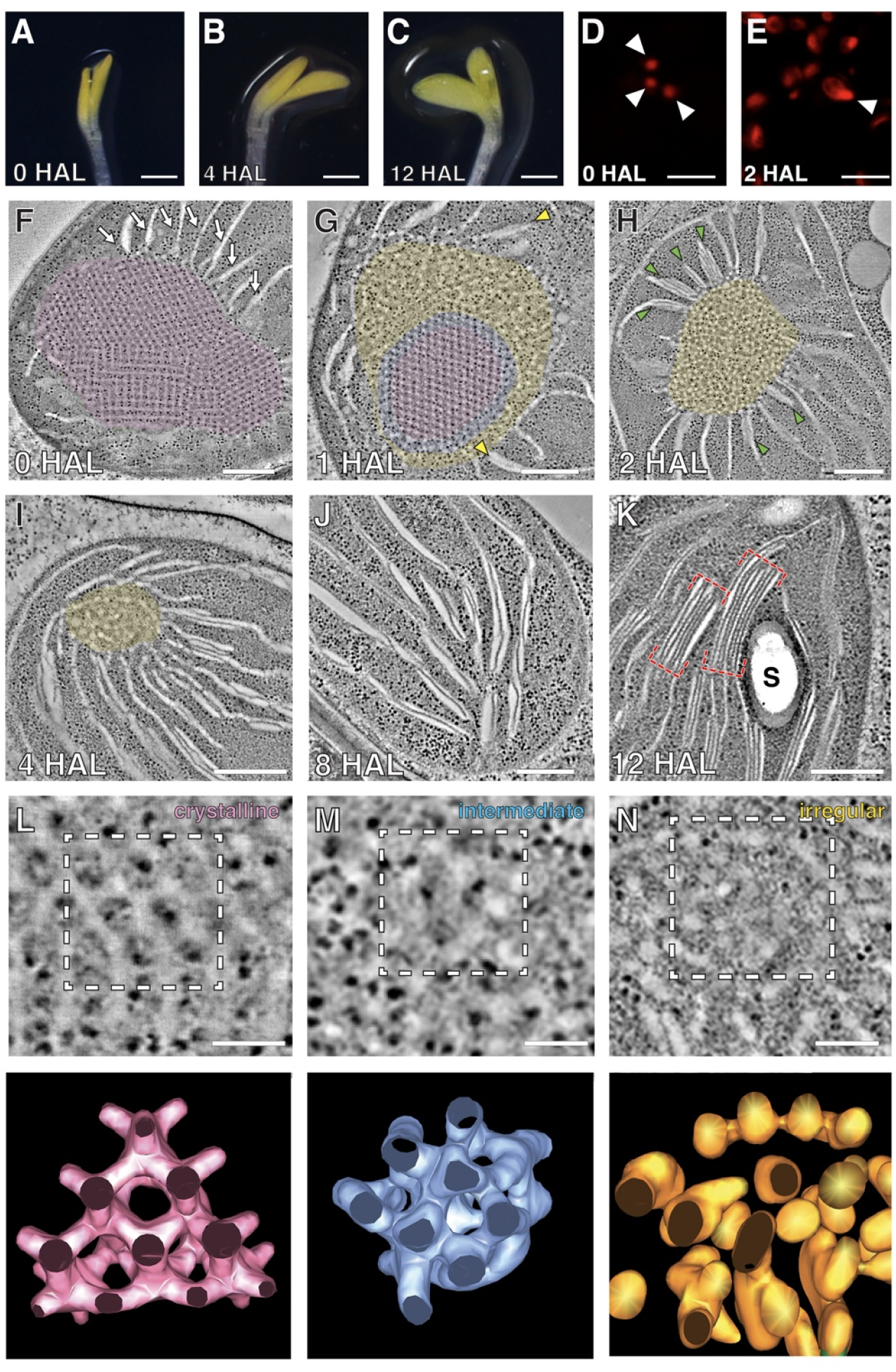
PLB degeneration and thylakoid assembly in de-etiolating *Arabidopsis* cotyledons. (*A-C*) 7-day-old etiolated *Arabidopsis* cotyledons at *A)* 0 HAL, *B)* 4 HAL, and *C)* 12 HAL. *(D-E)* Confocal laser scanning micrographs showing Pchlide/chlorophyll autofluorescence at *D)* 0 HAL and *E)* 2 HAL. Arrowheads indicate PLBs. *(F-K)* STET slice images of plastids at *F)* 0 HAL, *G)* 1 HAL, *H)* 2 HAL, *I)* 4 HAL,*J)* 8 HAL, and *K)* 12 HAL. The crystalline, irregular, and intermediate zones in PLBs are highlighted in magenta, yellow, and blue, respectively, in *(F)-(I)* Yellow arrowheads in *(G)* mark grana stacks of two layers. Green arrowheads in *(H)* indicate PLB-associated grana stacks. Grana stacks in *(K)* are denoted with red brackets. S: starch particle. Scale bars = 300 nm. (*L-N*) High magnification tomographic slice images of the PLB lattice (crystalline) at 0 HAL *(L)*, PLB tubules of the intermediate zone at 1 HAL *(M)* and PLB tubules of the irregular zone at 1 HAL *(N)* Scale bars = 150 nm. Lower panels show 3D surface models of the PLB membranes demarcated with dashed squares in upper images.

PLBs shrank quickly and lost their crystalline regularity by 2 HAL (Fig. 1G-H and S1G). Grana stacks of two layers appeared in the vicinity of PLB as early as 1 HAL (Fig. 1G), and the number of disks had increased in the PLB-associated grana stacks at 2 HAL (Fig. 1H). PLBs were almost degraded at 4 HAL and had disappeared completely in 8 HAL samples (Fig. 1I-J and S1H-I). Grana stacks and stroma thylakoids were evident in cotyledon cells at 12 HAL, and the chloroplasts had starch particles (Fig. 1K and S1J). Chloroplasts at 12 HAL were approximately 25% larger than etioplasts at 0 HAL (Fig. S1K).

Loss of the crystalline architecture began at the PLB surface (Fig. 1G). During the loss of the crystalline architecture, the inner core retained the lattice structure, and tubules in the periphery were highly convoluted (Fig. 1G). In between the crystalline core and the irregular periphery was a narrow band in which the lattice arrangement was compromised (Fig. 1G). No crystalline symmetry was discerned in PLBs at 2 HAL, but grana stacks were detected at the PLB surface (Fig. 1H), suggesting that grana-forming super-complexes containing LHCII are assembled where chlorophyll molecules are produced from protochlorophyllides in the PLB.

We generated 3D surface models of PLB tubules in the periphery of PLBs and in the intermediate zone of PLBs from 0 and 1 HAL images. PLBs before illumination consisted of tetravalent nodes (Fig. 1L), and the average width of the tubules was calculated to be 24.0 nm (n = 46, SD = 3.11). In the intermediate zone at 1 HAL, tubular nodes were displaced, obscuring the hexagonal pattern (Fig. 1M). The peripheral tubules were highly disordered and had varying thicknesses (Fig. 1N).

### Skeleton models of PLBs

We converted PLBs in our tomograms into 3D skeleton models consisting of lines and nodes using the MATLAB software package (Fig. S2). Hexagonal and square lattices were discerned in the 0 HAL skeleton model (Fig. 2A-B). Among crystallographic symmetries associated with PLBs, the diamond cubic symmetry matched our PLB skeleton model. We were able to capture views of the models corresponding to the Miller index planes of the diamond cubic symmetry (Fig. 2C-E). The hexagonal periodic patterns conformed to the (1,1,1) or (1,1,0) planes, whereas the square lattice matched the (1,0,0) plane. The diamond cubic unit cell size averaged to 65.5 nm (n = 61, SD = 3.71) when measured from nodes in the (1,1,0) or (1,0,0) planes.

**Figure 2.**
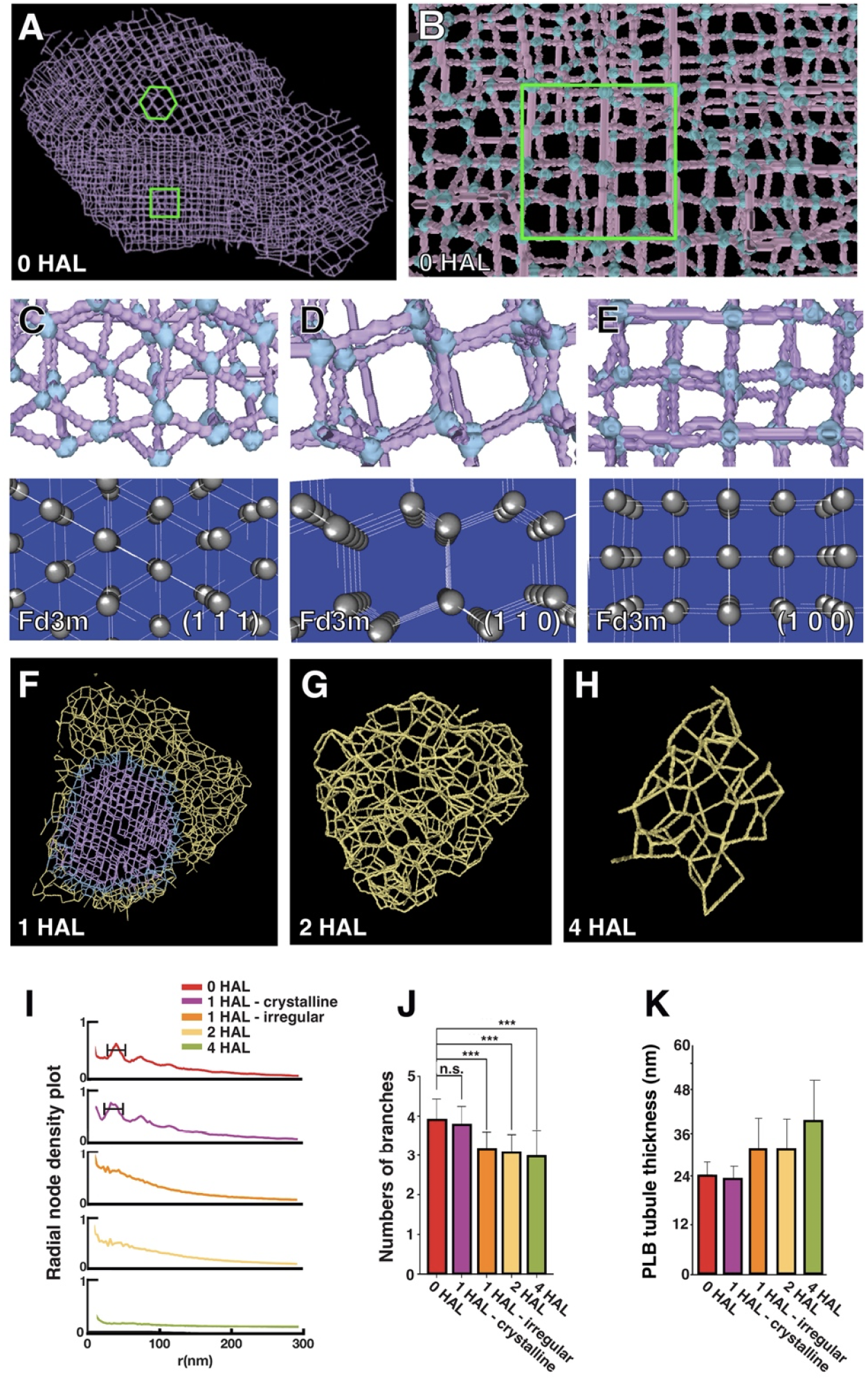
The crystalline structure of *Arabidopsis* PLB and its decay during de-etiolation. *(A)* A skeleton model of the PLB in Fig. 1F. Regions exhibiting hexagonal or square lattice patterns are marked in green. (B) A higher magnification view of the skeleton model shown in panel A. Nodes are highlighted in light blue. The region exhibiting a square lattice pattern is marked in green. (C-E) Projection views of select regions (upper panels), and lattice planes of the space group Fd3m (cubic diamond crystal structure) and their Miller indices, (1,1,1), (1,1,0), and (1,0,0) of the PLB skeleton model in panel B (bottom panels). Note that arrangements of PLB nodes and tubules match those of the cubic diamond lattices in all three planes. *(F-H)* Skeleton models of decaying PLBs at *F)* 1 HAL, G) 2 HAL, and *H)* 4 HAL. The models were generated from the tomograms in Fig. 1G, H, and I, respectively. Lines are color-coded to denote the crystalline, irregular, and intermediate zones in PLBs. (*I*) Radial density plots of branching nodes at four timepoints of de-etiolation. *(J)* The average numbers of branches at each node in 0 HAL, 1 HAL crystalline, 1 HAL irregular, 2 HAL, and 4 HAL PLBs. *(K)* The average diameters of tubules in 0 HAL, 1 HAL crystalline, 1 HAL irregular, 2 HAL, and 4 HAL PLBs.

From the skeleton models from 1, 2, and 4 HAL PLBs (Fig. 2F-H), we calculated radial densities of nodes and branching numbers per node at each time point. The node density plot had a peak from 30 nm to 70 nm in 0 HAL and 1 HAL crystalline PLBs, indicating a regular spacing between nodes (Fig. 2I). The peak was not present in irregular region of PLBs at 1 HAL. In agreement with the tetravalent units seen in 3D models (Fig. 1M), each node had four branches in 0 HAL and 1 HAL crystalline PLBs (Fig. 2J). Numbers of branches decreased as PLBs were degraded in later time points (Fig. 2K) and the reduction was accompanied by an increase in tubule thickness (Fig. 2L).

### Assembly of lamellar thylakoids and grana stacks on the PLB surface

3D tomographic models of the PLB-thylakoid interface at 1, 2, and 4 HAL were generated to examine how PLB tubules give rise to pre-granal thylakoids and how they turn into grana stacks. The irregular tubules observed in the PLB periphery at 1 HAL became interwoven and smoothened to constitute fenestrated membrane sheets at later time points. They consolidated into pre-granal thylakoids as their openings shrank and disappeared (Fig. 3A-C). Thylakoids in the immediate vicinity of PLBs were fenestrated at 2 HAL, indicating that PLB tubules continued to fuse (Fig. 3D-E).

**Figure 3.**
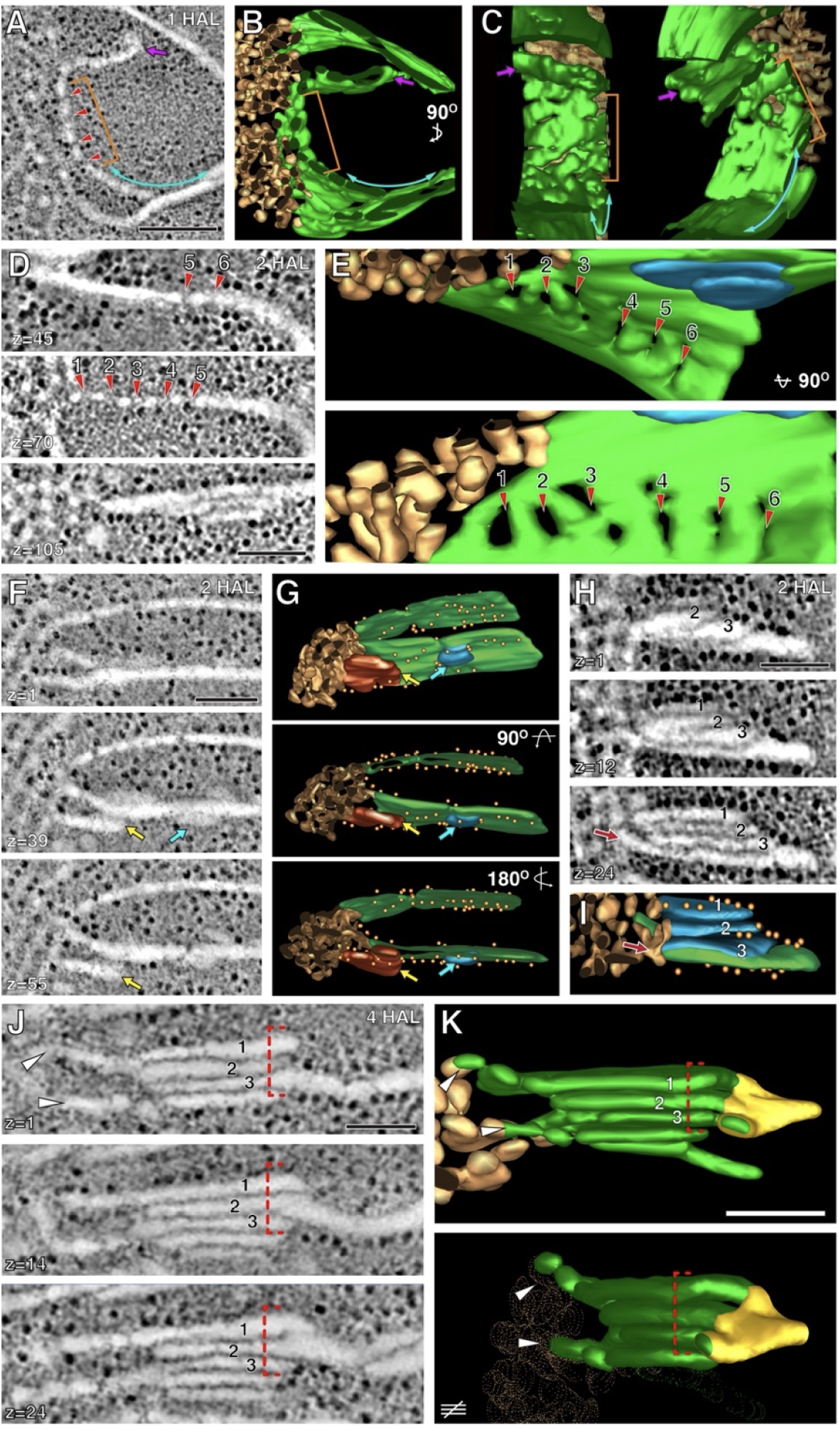
PLB to pre-granal thylakoid transition and grana formation from pre-granal thylakoids. *(A-E)* Tomographic slice images (panels A and 0) and 30 models of fenestrated sheets on the PLBs at 1 HAL. The membrane in panels 0-E is more consolidated than the sheet shown in panel A. Holes indicated by the red arrowheads. *(F-G)* High-magnification images of two pre-granal thylakoids connected to PLB periphery and their 30 models. Buds on pre-granal thylakoids are indicated by blue arrows. Connections with nascent pre-granal thylakoid are indicated by yellow arrows. *(H-I)* Image and 30 model of a grana-forming stack consisting of four layers at the margin of a decaying PLB. Three small lamellae (blue) derived from the irregular PLB margin are on a grana-forming thylakoid (green). They are interconnected via their edges. *(J-K)* Images and 30 model of a grana-stroma thylakoid that appears more mature than the grana-forming stack in panels F-H. The grana layers are connected to each other through membrane folds (red brackets). The model in the lower panel is a view after 90^°^ rotation. The helical arranged stroma thylakoids (yellow) interconnected the new grana stack. Scale bars = 100 nm.

Stacked thylakoids appeared on the PLB surface at 2 HAL where two pre-granal thylakoids were laterally fastened to each other or grana-forming buds emerged (Fig. 3F-G). Three or four-layered grana were also observed when thylakoid sheets overlap or fold repeatedly. The acquisition of new layers did not seem to occur in an orderly fashion (Fig. 3H-I). Their diverse membrane configurations were similar to those of pro-granal stacks in young chloroplasts of germinating cotyledon cells (Liang et al. 2018). Grana stacks displaced from PLBs were frequently observed at 4 HAL (Fig. 3G-H). They had five or six disks interconnected by helical stroma thylakoids as typical grana stacks do.

To correlate expression levels of thylakoid membrane proteins with thylakoid assembly, we performed RNA-seq analysis of de-etiolating cotyledon samples isolated at 0, 1, 2, 4, 8, and 12 HAL. Most components of the photosystems and light harvesting complexes were upregulated at 2 HAL when grana stacks appeared on the PLB surface (Fig. S3A). Levels of *Lhcb*, which encodes a protein necessary for thylakoid stacking, sharply increased between 1 HAL and 2 HAL (Fig. S3B). The mRNAs encoding PSII components and LHCB were generally more abundant than mRNAs encoding PSI constituents and LHCA before 2 HAL.

### CURT1A localizes to the nascent pre-granal thylakoids emerging from PLBs and grana stacks

Pre-granal thylakoids and grana stacks emerging from the PLB surface had membranes sharply curved membrane toward the stroma (Fig. 3). This led us to hypothesize that CURT1 family proteins, which stabilize the grana margin, are involved in stack assembly from PLB tubules (Armbruster et al., 2013; Pribil et al., 2014). Among the four *CURT1* family genes, *curt1a* (AT4G01150) were more abundant than other members of the family in de-etiolating Arabidopsis cotyledons (Fig. S3C). We generated transgenic *Arabidopsis* lines expressing a CURT1A-GFP fusion protein under control of its native promoter to monitor its localization during de-etiolation. The fusion protein construct rescued the granum assembly defects of *curt1a-1* mutant cotyledons, indicating that the fusion protein is functional (Fig. S4 and S5). At 0 HAL, GFP fluorescence colocalized with PLB autofluorescence, although this colocalization was not complete; some PLBs had a GFP halo or GFP-positive puncta around them (Fig. 4A). In 2 and 4 HAL cells, PLB became smaller and chlorophyll autofluorescence spread over de-etiolating chloroplasts, and CURT1A-GFP formed foci on PLBs in 2 HAL chloroplasts (Fig. 4B). As thylakoids permeated the stroma, small GFP spots scattered to multiple locations in 4 HAL chloroplasts (Fig. 4C). CURT1A-GFP at 4 and 8 HAL were round or rod-shaped (Fig. 4D). We verified the association of CURT1A with PLBs with immunogold labeling. CURT1A-specific gold particles associated most frequently with the PLB surface where new thylakoid elements arise at 0 and 2 HAL (Fig. 4E-G and J). As PLBs were being consumed at 4 and 8 HAL, the majority of CURT1A relocated to thylakoids, binding to grana stacks in the stroma (Fig. 4H-L and J).

**Figure 4.**
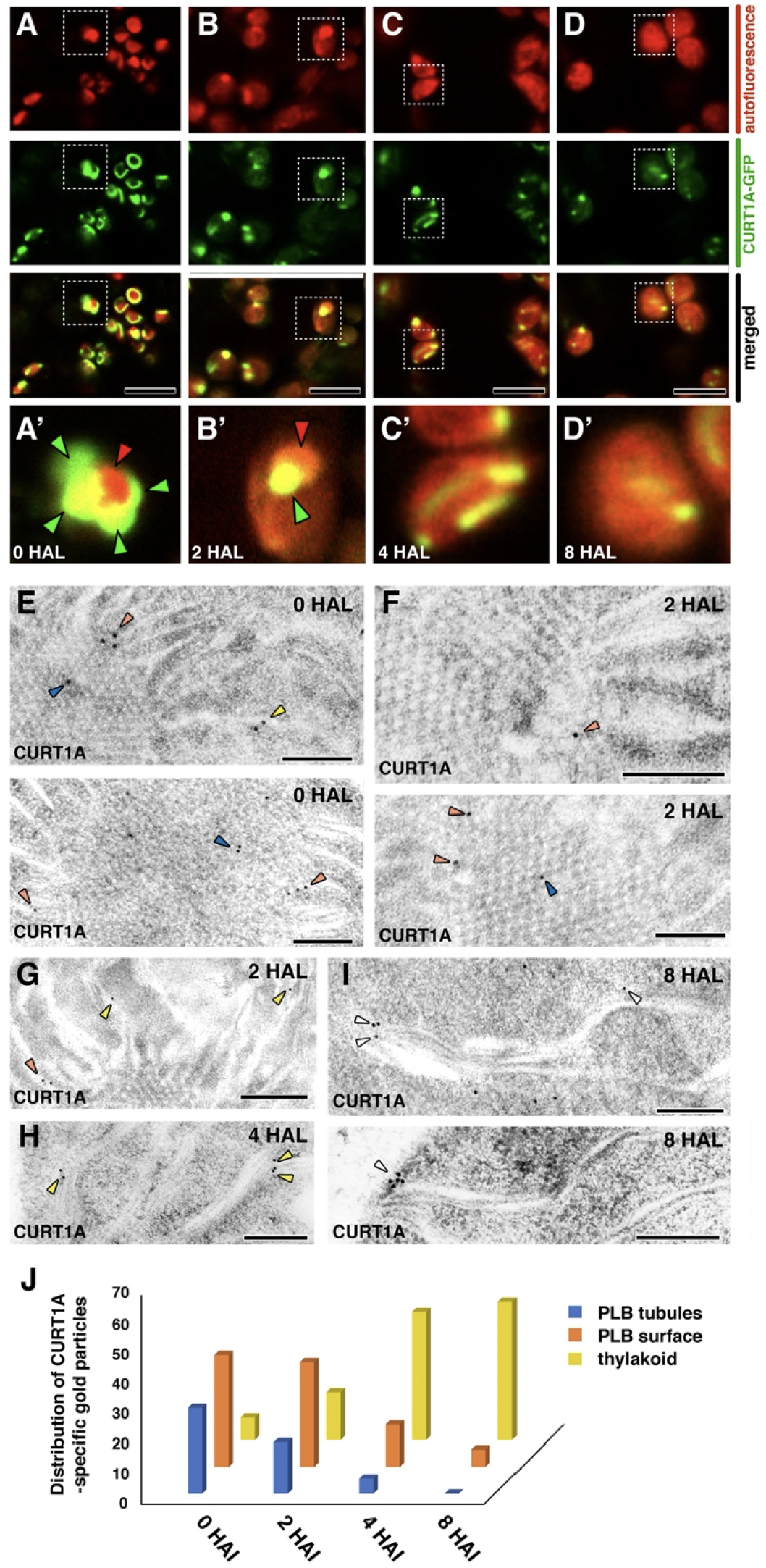
Localization of CURT1A proteins during de-etiolation. *(A-D)* Confocal laser scanning micrographs showing CURT1 A-GFP distribution at A) 0 HAL, B) 2 HAL, C) 4 HAL, and D) 8 HAL. Autofluorescence from Pchlide/chlorophyll, fluorescence from CURT1A-GFP, and merged panels are shown in each column. Panels A’-O’ are high-magnification micrographs of regions indicated with squares in panels A-D. In A’ and B’, PLBs and CURT1A-GFP puncta are indicated with red and green arrowheads. Scale bars = 8 *µm. (E-I)* lmmunogold labeling localization of CURT1A in *Arabidopsis* plastids at E) 0 HAL, F) 1 HAL, G) 2 HAL, H) 4 HAL, and I) 8 HAL. Gold particles located in PLBs, periphery of PLBs, and thylakoids are marked with blue, orange and yellow arrowheads, respectively. Scale bars= 200 nm. (J) Histogram showing CURT1A-specific gold particle distribution in *Arabidopsis* plastids at 0 HAL, 1 HAL, 2 HAL, and 4 HAL.

### Assembly of pre-granal thylakoids and grana stacks is abnormal in *curt1a* mutant cotyledons

To test whether CURT1A is required for the pre-granal thylakoid assembly and grana formation, we isolated *curt1a* T-DNA inserted mutant lines (Fig. S4 and S5). Etioplasts in 0 HAL *curt1a-1* (SALK_030000) cotyledons appeared normal. However, at 1 HAL, thylakoids on the PLB surface were swollen in mutant cotyledon cells in contrast to the flat pre-granal thylakoids in wild-type cotyledon cells (Fig. 5A-B). The bloated thylakoids failed form grana stacks. Instead, round thylakoids accumulated around PLBs at 2 and 4 HAL in the mutant (Fig. 5C-D). Despite the lack of grana stack formation, PLBs shrank and stroma thylakoids continued to expand (Fig. 5E). Chloroplasts had extremely extended stacks made of two or three layers in *curt1a-1* mutant cotyledon cells at 12 HAL (Fig. 5F). Another T-DNA mutant allele of *curt1a* (*curt1a-2*, GK-805B04) also had swollen thylakoids and lacked grana stacks, and the phenotype was rescued upon transformation with the CURT1A-GFP construct (Fig. S5G-K).

**Figure 5.**
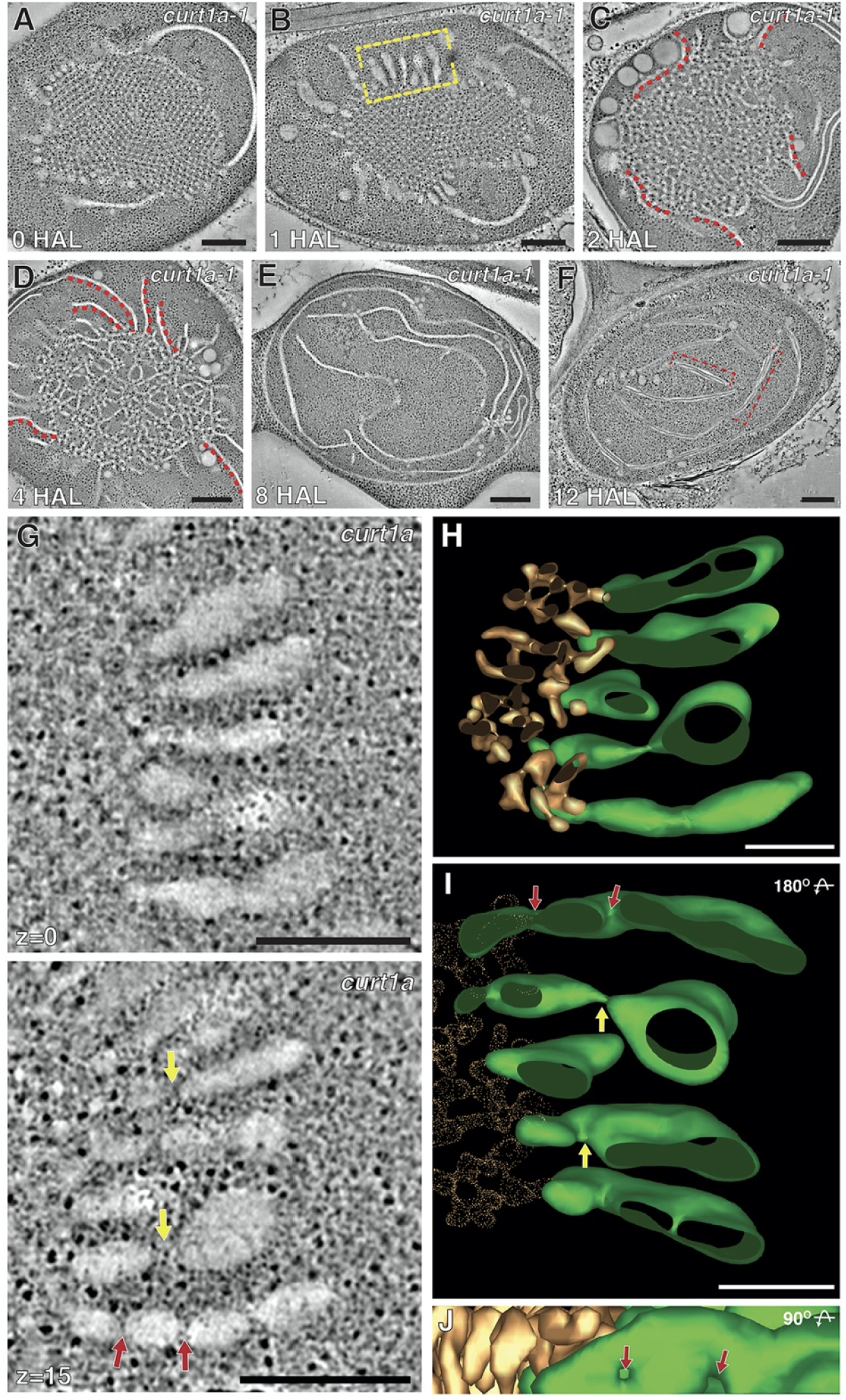
Plastid de-etiolation is altered in the *curt1a* mutant. *(A-F)* STEM tomography slices of *curt1 a-1* plastids at A) 0 HAL, B) 1 HAL, C) 2 HAL, 0) 4 HAL, E) 8 HAL, and F) 12 HAL. Red dots in *(CJ* and *(DJ* label pre-granal thylakoids associated with PLBs but no grana stacks arise from them. Scale bars= 300 nm. (G) A tomographic slice image of the thylakoids connected to PLBs in the yellow bracket in panel B. Fenestrations in pre-granal thylakoids are indicated by red and yellow arrows. Scale bars= 200 nm. *(H-J’J* 30 model of the image in panel G with thylakoids in green and PLBs in gold. The 30 model in panel I is rotated 180^°^ relative to that in panel H and that in panel J is rotated an additional 90^°^. Fenestrations are indicated by yellow and red arrows. Scale bars= 100 nm.

### The cubic crystalline lattice is interrupted in *curt1c* mutant PLBs

As transcripts from *curt1b* (AT2G46820) and *curt1c* (AT1G52220) accumulated in de-etiolating cotyledon cells, we examined T-DNA mutant lines in which *curt1b* or *curt1c* was inactivated (Figs. 6 and S6). We noticed that PLBs of *curt1c*-*1* (SALK_023574) cotyledons often had irregularities (Fig. 6 A, B, and J). Pores as large as 400 nm (Fig. 6A) or areas where PLB tubules were disorganized (Fig. 6B) were seen *curt1c-1* PLBs. However, they did not have swollen thylakoids around PLBs and grana stacks developed in association with degrading PLBs at 4 HAL (Fig. 6D-E). When we expressed CURT1C-GFP from its native promoter in the *curt1c-1* background, the defects in PLBs disappeared (Fig. 6C and J).

**Figure 6.**
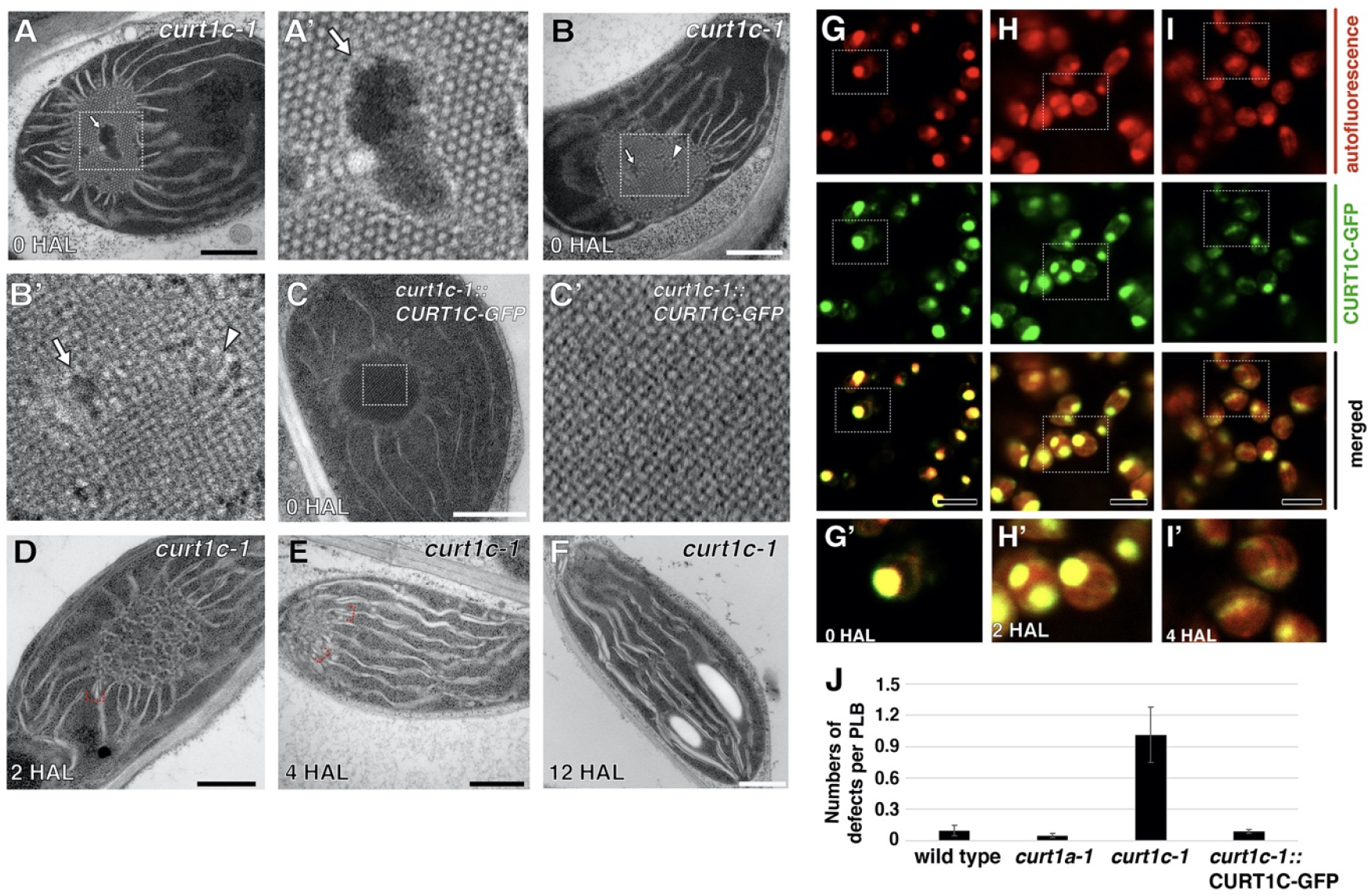
PLBs in *curt1c* mutant etioplasts are abnormal. *(A-BJ* TEM micrographs of PLBs in *curt1c-1* etioplasts at O HAL. Holes and disordred tubules are marked with arrows and arrowheads, respectively. (CJTEM micrograph of an etioplast in *curt1c-1* expressing CURT1C-GFP at O HAL. A’-C’ are magnified views of PLBs inside the rectangles in panels A-C. *(D-F)* TEM micrographs of *curt1c-1* plastids at D) 2 HAL, E) 4 HAL, and F) 12 HAL. Grana stacks associated with PLBs are denoted with brackets in panels D and E. *(G-H)* Confocal laser scanning micrographs showing CURT1C-GFP distribution at G) 0 HAL, H) 2 HAL, and I) 4 HAL. Autofluorescence from Pchlide/chlorophyll, fluorescence from CURT1C-GFP, and merged panels are shown in each column. Panels G’-I’ are high-magnification micrographs of regions denoted with squares in panels G-1. *(J)* Frequencies of PLB irregularities. The histogram was produced after analyzing ∼30 etioplasts from at least three samples for each genotype. Most PLBs had pores or abnormal tubules in *curt1c-1*.

CURT1C-GFP expressed by the CURT1C promoter overlapped almost completely with PLB autofluorescence at 0 HAL and shrank together with PLBs (Fig. 6E-F). This contrasted with CURT1A-GFP which enclosed PLBs or constituted foci around PLBs (Fig. 6G-H). In 4 HAL plastids where PLBs have been mostly depleted, GFP-positive spots were scattered over thylakoids in a manner similar to that of CURT1A-GFP (Fig. 6I). PLBs, their degradation, and thylakoid development around PBLs appeared normal in *curt1b-1* (WiscDsLoxHs047_09D) cotyledon cells (Figs. S6). However, in all alleles of *curt1a, 1b*, and *1c*, grana stacks had fewer layers and were wider than those in wild-type chloroplasts at 12 HAL (Figs. 1, 5, 6, and S6).

## Discussion

We have determined the crystalline structure of *Arabidopsis* PLBs in high-pressure frozen etioplast samples to be zinc blende type. This result agrees with a small angle X-ray diffraction study of isolated maize PLBs (Selstam et al., 2007). We did not find any evidence in our 3D models for wurtzite-type lattices; this lattice was detected in runner bean by Kowalewska et al. (2016). The wurtzite lattice was seen at the boundary between zinc blende crystal domains in squash etioplasts (Murakami et al., 1985). The zinc blende lattice is a center-closest packed crystal system with a repeating unit of three layers, whereas the wurtzite lattice is a hexagonal closest packed system with a repeating unit of two layers (Cotton et al., 1995). We prepared approximately 300-nm thick sections that enclose more than four layers within PLBs and examined them with STET. Due to the dark staining of stroma in cryofixed samples, it was impossible to acquire tomograms in the brightfield mode with signal-to-noise levels sufficient for automatic segmentation. It was crucial to employ STET to enhance the membrane contrast in order to establish the crystalline structure of PLBs.

The LPOR–Pchlide–NADPH ternary complex binds to the lipid bilayer to produce membrane tubules *in vitro*, and the complex breaks apart upon illumination to release raw materials necessary for constructing photosynthetic thylakoids (Nguyen et al., 2021). We observed collapse of crystalline order from the peripheral PLBs at 1 HAL, indicating that light activation of LPOR and subsequent Pchilde reduction began from the PLB exterior. The loss of crystalline architecture at 1 HAL was characterized by randomized internodal distances, reduced branching per nodes, and thickening of tubules. The tetrahedral branching points were dislocated in the intermediate zone in 1 HAL PLBs. Lying between the inner crystalline and outer irregular regions, the intermediate zone is likely the site of PLB lattices in which LPOR–Pchlide–NADPH ternary complexes have disassembled immediately after illumination.

One of the first events in the conversion of proplastids into chloroplasts is the formation of pre-granal thylakoids from tubule-vesicular prothylakoids (Liang et al., 2018). Our data indicate that at the PLB-stroma interface, pre-granal thylakoids develop from PLB tubules, and grana stacks arise from the pre-granal thylakoids (Fig. 7A). Localization of CURT1A, CURT1A-GFP, and *curt1a* mutant phenotypes provided evidence that CURT1A stabilize the curve membrane at the tip of pre-granal thylakoid emerging from PLBs. In *curt1a* mutant lines, we observed swollen thylakoids in association with degrading PLBs (Fig 7B). CURT1A-GFP complemented the *curt1a* phenotype, and the fusion protein localized to the PLB margin. We hypothesize that the sites in PLBs where CURT1A-GFP concentrated at 0 and 2 HAL correspond to sites from which new pre-granal thylakoids and grana stacks develop. No grana stacks developed in the vicinities of PLBs in *curt1a* mutants in agreement with the observed recruitment of CURT1A to the PLB periphery. All three *curt1* isotypes, *1a, 1b*, and *1c*, were transcriptionally active, and their gene products were detected in de-etiolating cotyledon specimens. The *curt1b* and *curt1c* mutant lines did not exhibit defects in the transition of PLBs into pre-granal thylakoids.

**Figure 7.**
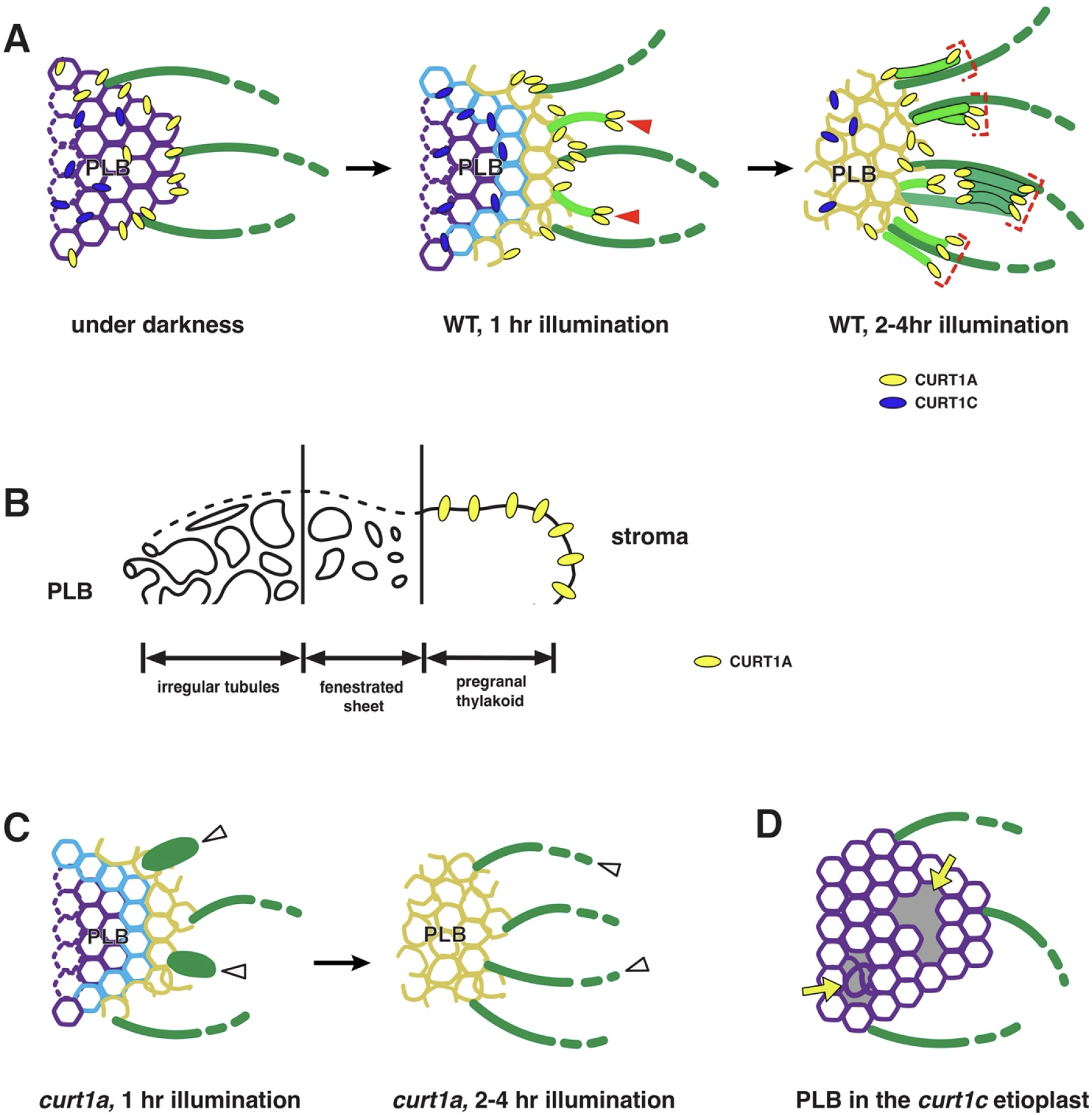
Schematic diagrams illustrating functions of CURT1A and CURT1C during de-etiolation. *(A)* Both CURT1 A and CURT1 Care localized in PLBs. Upon illumination, PLB decay occurs from the margin, and CURT1 A concentrates to the sites where new pre-granal thylakoids emerge (arrowheads) and grana stacks arise (brackets). CURT1 C does not exhibit such relocation dynamics. *(B)* Model of pre-granal thylakoid development from PLB tubules. The irregular region of degrading PLBs resemble a tubule-vesicular meshwork that gradually coalesces into a fenestrated sheet. CURT1 A bends the membrane at the edge of a nascent pre-granal thylakoid outgrowing from a PLB into the stroma. (C) Pre-granal thylakoids growing out from PLBs in *curt1a* are swollen (arrowheads) and they elongate without creating grana stacks. (0) PLBs lacking CURT1 C have pores and disorganized tubules (arrows).

The morphological gradient from tubules into fenestrated sheets, and further into mature pre-granal thylakoids (Fig. 7C) is reminiscent of cell plate maturation (Otegui et al., 2001; Otegui and Staehelin, 2004; Segui-Simarro et al., 2004). Vesicle fusion at the cell plate produces tubules that merge with each other to form fenestrated sheets. As the cell plate expands and matures, fenestrated sheets are consolidated into a new cell boundary (Samuels et al., 1995; Kang et al., 2003; Ahn et al., 2017). The membrane dynamics from tubules to planar thylakoids is completed in a miniature scale within several hundred nanometers from the PLB. However, the pre-granal thylakoid formation we observed was oriented backward from the cell plate assembly. Unlike the cell plate where vesicles and tubules are recruited to the growing end, sheet-like thylakoids outgrow from PLBs into the stroma.

PLBs in *curt1c* mutant cotyledons had large holes or disarrayed tubules, indicating that CURT1C is required for PLB assembly in darkness. When we expressed CURT1C-GFP from the native *curt1c* promoter, the defects were rescued, indicating a skotomorphogenic function of CURT1C (Fig. 7D). During de-etiolation, CURT1C-GFP spread uniformly over PLBs, and the signal faded together with PLB degradation. This GFP localization contrasted with that of CURT1A-GFP. The distinct phenotypes of *curt1a* and *curt1c* suggest that CURT1A selectively interacts with the machinery in the PLB periphery producing pre-granal thylakoids and grana stacks. It will require 3D electron microscopic analyses of etiolating plastids at multiple time points after seedling germination under darkness to characterize functions of CURT1C in PLB biogenesis.

In a recent publication, it was reported that chloroplast biogenesis from etioplasts and the PLB structure are affected in the *curt1abcd* quadruple mutant seedlings and that overexpression of CURT1A altered PLB morphology (Sandoval-Ibanez et al., 2021). We discovered that CURT1A and CURT1C contribute to the PLB-to-pre-granal thylakoid transformation and the development of semi-crystalline structure of PLBs, respectively. We were not able to confirm changes in PLBs after CURT1A overexpression as our CURT1A-GFP was expressed from the endogenous *curt1a* promoter. The defects we observed in *curt1a* and *1c* mutant lines are not in agreement with the observations documented by Sandoval-Ibanez et al. (2021) probably because *curt1abcd* specimens were processed by traditional protocols that involve fixation and dehydration at room temperature. It is generally accepted that membranous structures, including thylakoids, are preserved closer to their native states by cryofixation than by conventional fixation (Kiss et al., 1990; Kang, 2010; Nicolas et al., 2018; Otegui, 2021). Functional characterization of a plastid-targeted protein, FZL, provides an example. Inhibition of thylakoid membrane fusion in *fzl* mutant chloroplasts was discerned in high-pressure frozen samples (Liang et al., 2018) but not in chemically fixed samples in an earlier study (Gao et al., 2006).

## Methods

### Plant Materials and Growth Conditions

*Arabidopsis* Columbia (Col-0) and *curt1* seeds (NASC, http://arabidopsis.info/) were surface-sterilized and incubated in 4°C overnight. The seeds then were placed on 0.75% phytoagar Petri dishes supplemented with half-strength Murashige-Skoog salt (0.5 g/L, pH 5.8). The dishes were placed in a growth chamber (Cat No. MLR-352H-PB, Panasonic, Japan) at 22^°^C and were left to germinate and grow for 1 week under darkness. Samples were harvested after illumination with white fluorescent light at a photon flux intensity of 120 μmol m^-2^ s^-^ before dissection. The isolated cotyledon samples were analyzed by high-pressure freezing, RNA-seq, or confocal laser scanning microscopy.

### Chlorophyll extraction and measurement

Chlorophyll extraction and measurement were performed as described by Ma et al. (Ma et al., 2021). *Arabidopsis* seedlings were before incubated in 700 μL pre-heated DMSO in 65°C for 30 minutes and ground for centrifugation at 6000 rpm for 10 minutes. Another 300 μL pre-heated DMSO were added to the supernatant to make the final volume of 1 mL. The OD at 645 nm and 663 nm for chlorophyll a and chlorophyll b were measured with NanoDrop (Thermofisher). The Chlorophyll concentration was calculated with the Arnon’s equations. The experiment was repeated for three times and the quantification of chlorophyll concentration was performed using Microsoft Excel 2016, the graphs were made by Prism8 (GraphPad Software).

### Generation of CURT1A-GFP, CURT1C-GFP lines in their respective mutant backgrounds

The genomic fragment of *CURT1A* (AT4G01150) and *CURT1C* (AT1G52220) including ∼2 kb promoter region was amplified and inserted into a binary vector pBI121. The last exons of the genes were translationally fused with the GFP in the vector. *curt1a-1* and *curt1c-1* plants were transformed with the CURT1A-GFP and CURT1C-GFP constructs, respectively by floral dip method with the Agrobacterium tumefaciens strain GV3101(Zhang et al., 2006). Transgenic seedling (T1) were selected by kanamycin containing 1/2MS + 0.8% agar (w/v). Seedlings (T2 generation) were tested for GFP expression with immunoblot analysis (anti-GFP antibody, 1:2500 dilution, Abcam Cat# av290) and observed under Leica TCS SP8 Confocal Microscope System. All the primers were from Integrated DNA Technologies, and the genomic fragments were amplified with iProof high-fidelity DNA polymerase (Bio-Rad). The primer sequences for the GFP cloning are in Supplemental Table 2.

### High-pressure freezing (HPF), sample processing, and TEM

High-pressure freezing, freeze substitution, resin embedding, and ultramicrotomy were performed as described in Kang (Kang, 2010). Seedlings were examined with a Canon EOS M50 Digital Camera equipped with fluorescence illumination to remove abnormal cotyledons before freezing. Frozen samples were freeze-substituted in anhydrous acetone with 1% OsO4 at -80°C for 24 hr. Excess OsO4 was removed at -80^°^C by rinsing with precooled acetone. After being slowly warmed up to room temperature over 60 h, samples were separated from planchettes and embedded in Embed-812 resin (Electron Microscopy Sciences; catalog no. 14120). 80 nm thick sections of each time point were prepared with ultramicrotomy and then were examined with a Hitachi 7400 TEM (Hitachi-High Technologies) operated at 80 kV.

### Dual-axis scanning Transmission Electron Tomography (STEM), Tomogram Reconstruction, Modeling, and Measuring Morphometric Parameters

300 nm thick sections were collected on formvar-coated copper slot grids (Electron Microscopy Sciences; catalog no. GS2010-Cu) and stained with 2% uranyl acetate in 70% methanol followed by Reynold’s lead citrate (Mai et al., 2019). Tilt series from ±54° at 1.5° intervals in scanning transmission electron microscopy (STEM) mode were collected with a 200-kV Tecnai F20 intermediate voltage electron microscopy (Thermo-Fischer, USA). FEI STEM Tomography interface was used to collect two tilt series around the orthogonal axe as described in Kang (2016) and membrane surface models were prepared according to (Mai and Kang, 2017).

### Immunoblot Analysis and Immunogold Labeling

Protein samples were extracted from seedlings at 0 h, 1 h, 2 h, 4 h, 8 h and 12 h of light treatment after being pulverized in liquid nitrogen. SDS-PAGE and immunoblot were performed as described by Liang et al. (2018) and Lee et al. (Lee et al., 2013). The experiment was repeated for three times with total protein extracts from three independent sets of cotyledon samples. For immunogold labeling, thin sections (80 nm thick) of HM20 embedded samples at each time point were prepared by ultramicrotomy, the following immunodetection of gold particles were performed according to the protocol explained in Wang et al. (Wang et al., 2017). Antibodies for CURT1A (AS08 316), CURT1B (AS19 4289), CURT1C (AS19 4287) and PORA (AS05 067) were purchased from Agrisera. Anti-PBA1 antibody (ab98861) was purchased from Abcam.

### Transcriptomic Analyses

RNA samples were isolated from seedlings at each time point with 3 biological replicates using Qiagen Plant RNA extraction kit (Qiagen; catalog no. 74904). A total of 18 cDNA libraries were prepared following the standard BGISEQ-500 RNA sample preparation protocol and sequenced by the DNBseq platform (BGI, Shenzhen, China). Raw reads were filtered by SOAPnuke software, about 23.23 m clean reads for each sample were obtained in FASTQ format. The transcript expression level was then calculated and normalized to FPKM using RSEM software. The heat maps and the line charts were generated with R Studio (version 1.1.383) as described previously (Liang et al., 2018). FPKM values for CURT1 family genes and Lhcb were calculated to evaluate their expression level.

### Generation of skeleton models from PLB tubules

PLB membranes were first segmented using the 3D Orientation Field Transform tool (https://arxiv.org/abs/2010.01453). Skeletons were generated from the segmented membrane tubules by performing a medial axis transform (also known as ‘skeletonisation’) with an in-built MATLAB algorithm. Each skeleton element was converted into an undirected adjacency matrix carrying node coordinates using the Skel2Graph3D algorithm developed by (Kollmannsberger et al., 2017). The Bresenham’s line algorithm was used to connect node pairs with a straight line (https://arxiv.org/abs/2010.01453). The MATLAB adaptation of the Bresenham’s line algorithm iptui.intline() was modified for the purpose.

### Analysis of computer generated PLB skeleton models

The radial distribution function was computed by first plotting a histogram of the distances *r* between all the nodes in the skeleton, then the binned number was divided by *4πr*^*2*^. A curve approximating the histogram was used to generate probability plots against radial distances. Numbers of branches were counted from skeleton models at each time points. A distance transform with an in-built MATLAB algorithm was used on the binary segmented PLB tomograms to estimate PLB tubule thicknesses. The skeleton of the original segmentation was then introduced as a mask to select voxels around the central axes of PLB tubules. Approximate radii of PLB tubules were calculated from sizes of the voxels. The radii values were doubled to acquire diameters that correspond to tubular thicknesses. As we have calculated diameters from numerous voxels along auto-segmented PLB tubules, we were able to acquire low p-values (high degrees of confidence) in the pairwise comparisons.

### Accession Numbers

The RNA-seq data have been deposited in NCBI Sequence Read Archive under accession number GSE189497.

## Supplemental Data

The following material is available in the online version of this article.

Supplemental Figure 1. Etioplast-chloroplast transition in de-etiolating *Arabidopsis* cotyledons

Supplemental Figure 2. Generating skeleton models from PLB tubules in tomograms.

Supplemental Figure 3. Analyses of transcripts in de-etiolating *Arabidopsis* cotyledons that encode proteins associated with photosynthesis.

Supplemental Figure 4. Characterization of *curt1* T-DNA inserted mutant lines.

Supplemental Figure 5. The abnormal thylakoid assembly phenotype reproduced in the *curt1a-2* (GK-805B04) allele and rescue of *curt1a* defects by expression of CURT1A-GFP

Supplemental Figure 6. Etioplast-to-chloroplast differentiation in *curt1b-1* cotyledons.

Supplemental Table 1. Statistics of the three rounds of RNA-seq experiments at the six time points.

Supplemental Table 2. Primer sequences for genotyping or molecular cloning

Supplemental dataset 1. The skeletal model file of a PLB shown in Figure 3.

## Acknowledgements

This work was supported by the Hong Kong Research Grant Council (GRF14121019, 14113921, AoE/M-05/12, C4002-17G) and Chinese University of Hong Kong (Direct Grants).

## Author Contributions

B-H. K. and Z.L. designed the research. Z.L., W.T.Y., K.K.M., J.M. Z.L. and Y-L. F. C. performed the experiments. All authors analyzed the data. B-H. K. and Z.L. wrote the article.

## Competing Financial Interest Statement

The authors declare no competing financial interest.

